# Crosstalk between Fat Mass and Obesity-related (FTO) and multiple WNT signaling pathways

**DOI:** 10.1101/2021.05.20.444911

**Authors:** Hyunjoon Kim, Soohyun Jang, Young-suk Lee

## Abstract

Fat Mass and Obesity-related (FTO) gene is associated with a diverse set of human diseases. Yet, the functional landscape of FTO remains largely unknown, most likely owing to its wide range of mechanistic roles and cell-type-specific targets. Here, we discover the intricate role of FTO in multiple WNT signaling pathways. Re-analyses of public data identified the bifurcation of canonical and noncanonical WNT pathways as the major role of FTO. In FTO-depleted cells, we find that the canonical WNT/β-Catenin signaling is inhibited in a non-cell autonomous manner via the upregulation of DKK1. Simultaneously, this upregulation of DKK1 promotes cell migration via activating the noncanonical WNT/PCP pathway. Unexpectedly, we also find that the canonical WNT/STOP signaling induces the accumulation of cytoplasmic FTO proteins. This subsequently leads to the stabilization of mRNAs via RNA demethylation, revealing a previously uncharacterized mode of WNT action in RNA regulation. Altogether, this study places the functional context of FTO at the branching point of multiple WNT signaling pathways which may explain the wide spectrum of FTO functions.

## INTRODUCTION

Fat mass and obesity-associated (FTO) gene is associated with body mass index (BMI), obesity risk and type 2 diabetes (Frayling et al., 2007; Sanghera et al., 2008). Loss-of-function mutations of FTO also cause severe growth retardation and reduced brain volume (Boissel et al., 2009; Caglayan et al., 2016; Daoud et al., 2016). FTO is also one of the six genes deleted in Fused toe (Ft) mouse mutant (Peters et al., 1999; van der Hoeven et al., 1994). Despite growing evidence that FTO is implicated in a broad range of human disease, the exact physiological role of FTO is largely unknown.

FTO is an alpha-ketoglutarate-dependent dioxygenase that demethylates mRNAs, tRNAs, and snRNAs to regulate RNA stability and function (Alarcon et al., 2015; Jia et al., 2011; Liu et al., 2015; Mauer et al., 2017; Wei et al., 2018). In particular, FTO demethylates and thereby stabilizes the mRNA of MYC, CEBPA, ASB2 and RARA leading to the enhanced oncogenic activity in certain leukemia cells (Su et al., 2018). FTO also plays a role in gluconeogenesis and thermogenesis by targeting the FOXO1 mRNA (Peng et al., 2019). During motile ciliogenesis, we discovered that FTO demethylates the exon 2 of FOXJ1 mRNA to enhance RNA stability (Kim et al., 2021). Much effort has been made to understand the molecular mechanisms of FTO, yet the cellular functions and upstream regulators of FTO that ultimately manifest its diverse roles warrants further investigation.

WNT signaling pathways regulate a variety of biological processes in development and tissue homeostasis (Logan and Nusse, 2004). In the canonical WNT/β-Catenin signaling pathway, the Wnt ligand (e.g. WNT3a) binds to the transmembrane receptor Frizzled (FZD) and low-density lipoprotein receptor-related 5/6 (LRP5/6) to stabilize β-Catenin proteins mainly by glycogen synthase kinase-3 (GSK3) inhibition (Piao et al., 2008; Taelman et al., 2010). Then, the β-Catenin proteins with TCF/LEF transcription factors activate gene expression of specific targets to regulate cell proliferation and differentiation (MacDonald et al., 2009). In the non-canonical WNT/PCP signaling pathway, Wnt ligands (e.g. WNT5A, WNT11) binds to the FZD receptor and co-receptors (e.g. ROR or Ryk) to activate small GTPases such as Rho, Rac and Cdc42 and the c-Jun N-terminal kinase (JNK) (Lai et al., 2009). In addition to the cytoskeleton reorganization by the small GTPases, JNK phosphorylates c-Jun and the activated c-Jun along with transcription factors ATF2 and ATF4 orchestrate an intricate transcriptional network to regulate cell motility and tissue polarity (Lai et al., 2009). The other β-Catenin-independent, non-canonical WNT/Ca2+ pathway induces the G-protein coupled receptor signaling cascade to increase intracellular calcium levels and regulate calcium/calmodulin-dependent protein kinases (Katoh, 2017; Niehrs, 2012). It is worth mentioning that canonical WNT signaling also leads to the cytoplasmic stabilization of many other GSK3-target proteins besides β-Catenin, termed the WNT/STOP pathway, thus remodeling the proteomic landscape (Acebron et al., 2014; Kim et al., 2015; Ploper et al., 2015).

In this study, we discovered a functional link between FTO and multiple WNT signaling pathways. Functional genomics analysis of FTO-depleted cells revealed specific downregulation of canonical WNT and upregulation of noncanonical WNT/PCP signalings. We further find that the canonical WNT/β-Catenin pathway is inhibited in FTO-depleted human cells. Among known WNT inhibitors, only DKK1 is upregulated after FTO-depletion and inhibits canonical WNT activity in a non-cell autonomous manner. This FTO-dependent DKK1 upregulation is also responsible for the enhanced cell migration of FTO-depleted cells by activating the noncanonical WNT/PCP pathway. Moreover, we report an unexpected role of the canonical WNT signaling pathway on mRNA stability via inhibiting GSK3-dependent FTO protein destabilization. Overall, our results unravel the crosstalk between FTO and WNT signaling pathways which may underlie the multifaceted complexity of FTO functions.

## RESULTS

### Geneset enrichment analysis hints at functional interactions between FTO and WNT signaling pathways

To investigate the functional role of FTO, we strategically re-analyzed a published FTO and ALKBH5 siRNA knockdown and RNA-Seq dataset (Mauer et al., 2017). Both FTO and ALKBH5 are the responsible enzymes for RNA N6-methyladenosine (m6A) demethylation (Jia et al., 2011; Zheng et al., 2013). m6A is the most abundant internal mRNA modification (Desrosiers et al., 1974), and thus FTO and ALKBH5 may generally regulate a common set of biological processes. To investigate the FTO-specific biological functions, we therefore utilized the ALKBH5 siRNA knockdown RNA-Seq dataset as a negative control for our geneset enrichment analysis. Specifically, we compared the geneset enrichment scores of differentially expressed genes in FTO-depleted cells against those in ALKBH5-depleted cells (Figures 1A and Figure 1-figure supplement 1A). Of note, negative enrichment scores indicate that genes of the set (e.g. Wnt signaling) are overall downregulated after siRNA treatment. Wikipathway geneset analysis (275 genesets) revealed negative enrichment of “Cytoplasmic Ribosomal Proteins (WP477)” and “Amino Acid metabolism (WP3925)” for both FTO- and ALKBH5-depleted cells (Figure 1A). Interestingly, “Wnt Signaling (WP428)”, “Mesodermal Commitment (WP2857)”, and “Endoderm Differentiation (WP2853)” were only downregulated in FTO-depleted cells (Figure 1A). Moreover, gene ontology enrichment analysis (3091 genesets) resulted in a negative enrichment of early developmental and cell proliferation-related biological processes specifically in FTO-depleted cells (Figure 1-figure supplement 1A). WNT signaling is one of the major signaling pathways governing cell proliferation and early developmental processes such as endomesodermal specification, indicating that FTO might be coupled with WNT signaling pathways.

**Figure 1.**
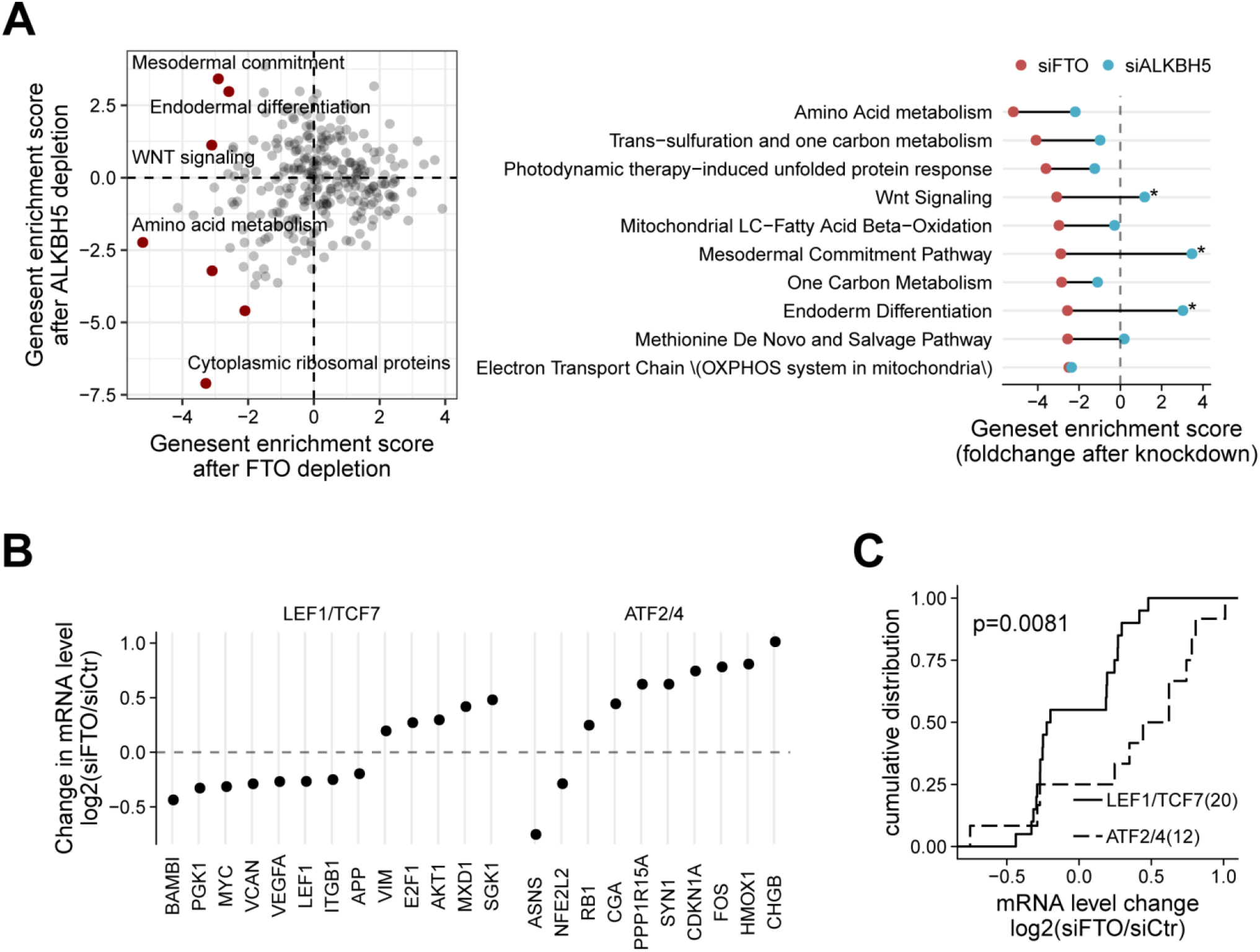
Systems approach unravels potential interaction of FTO with WNT signaling pathways. (A) Enrichment score of 275 wikipathway geneset pathways in FTO-depleted and ALKBH5-depleted cells. Notable pathways are marked in red (left). Top 10 wikipathway genesets with greater negative enrichment score in siFTO-depleted cells (red dots) than in siALKBH5-depleted cells (blue dots). Genesets with enrichment score difference greater than 4 are marked (right). (B) Regularized fold change after FTO knockdown of differentially expressed genes (p-value < 0.05) of LEF1/TCF7 targets and ATF2/ATF4 targets. (C) Cumulative plot of LEF1/TCF7 targets (solid) and ATF2/ATF4 targets (dashed) after FTO knockdown. The number of genes are shown in parenthesis. One-sided Kolmogorov-Smirnov test was used to test statistical significance.

To test whether this perturbation of WNT signaling at the pathway level directly impacts the transcriptional landscape, we examined the change in mRNA level of known transcriptional targets of canonical and noncanonical WNT signaling pathways (Figure 1B, p-value < 0.05). Targets of the canonical WNT/β-Catenin transcription factors (i.e., TCF7 and LEF1) including MYC, VEGFA, and LEF1 were downregulated in FTO-depleted cells. This result indicates that the observed WNT disruption in FTO-depleted cells leads to the inhibition of its targets, particularly that of the canonical WNT/β-Catenin signaling pathway. Unlike the canonical WNT targets, targets of WNT/PCP pathway transcription factors (i.e., ATF2 and ATF4) such as CDKN1A, FOS, and HMOX1 were upregulated, suggesting that FTO may inhibit the noncanonical WNT/PCP pathway. In fact, FTO-depleted cells exhibit a statistically significant bifurcation of gene expression between the TCF7/LEF1 targets and ATF2/ATF4 targets (Figure 1C, p-value = 0.0081). A similar trend was also observed in other human cells after FTO depletion (Figure 1-figure supplement 1B), hinting that this specific coupling between FTO and distinct WNT signaling pathways might be common in multiple cellular contexts.

### Loss of FTO inhibits the canonical WNT/β-Catenin signaling pathway in a non-cell autonomous manner by upregulating DKK1 mRNA expression

We employed the development system of *Xenopus laevis* embryos to examine the physiological impact of FTO. Knockdown of FTO in *Xenopus* embryos led to the anteriorization of neurula stage embryos and the impairment of neural crest and neuronal differentiation (Figure 2A). These defects altogether are reminiscent of canonical WNT inhibition (Kiecker and Niehrs, 2001; Kim et al., 2012), suggesting that FTO may regulate the canonical WNT/β-Catenin signaling pathway during early embryogenesis. Of note, this is consistent with a previous study which reported the inhibition of canonical WNT/β-Catenin signaling in *Fto* knockout mouse MEF and in FTO silenced HEK 293T cells (Osborn et al., 2014).

**Figure 2.**
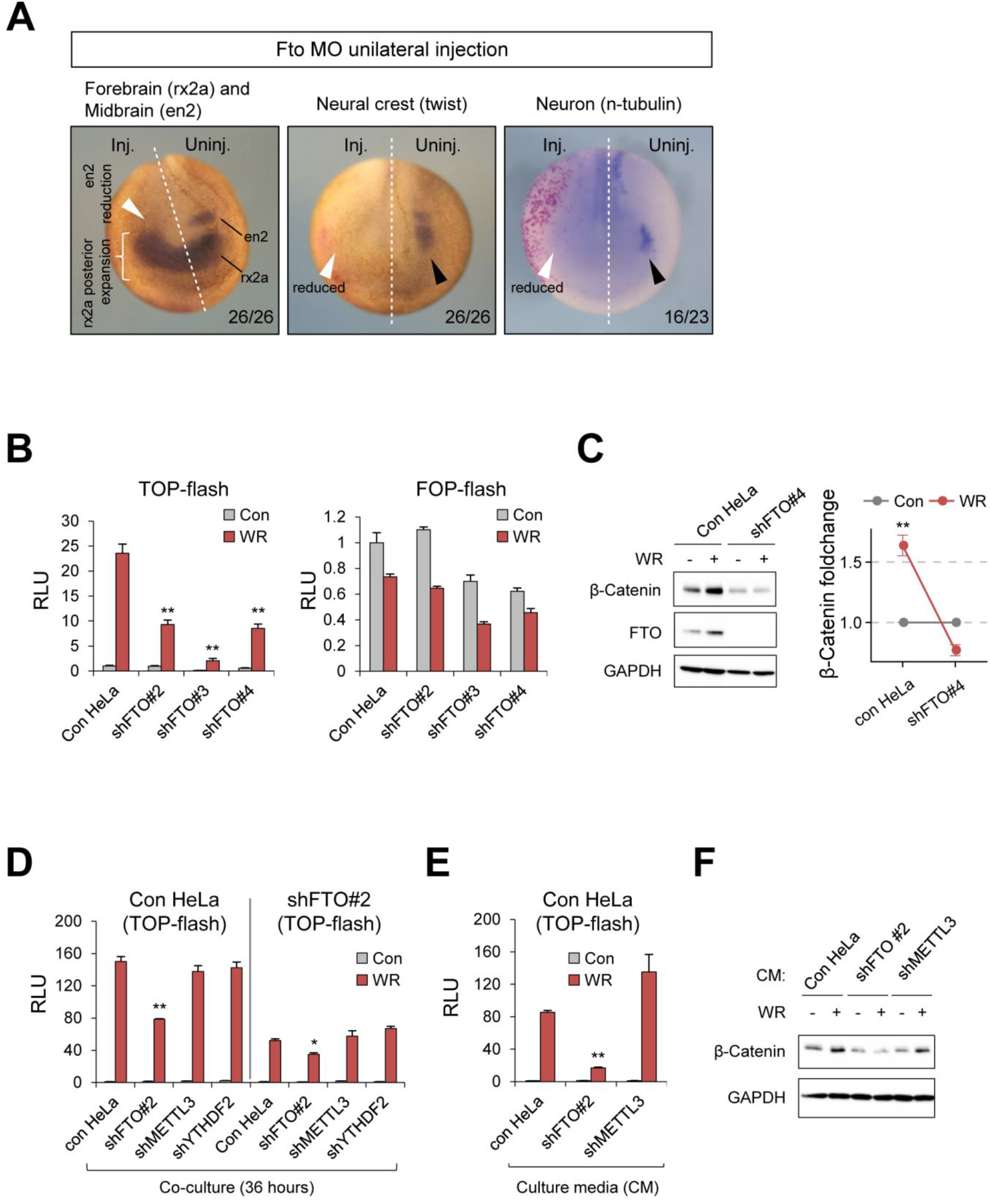
Loss of FTO supresses canonical Wnt/β-Catenin signaling. (A) In situ hybridization of neural patterning markers in *Xenopus* neurula embryos unilaterally injected with Fto MO. Forebrain marker rx2a was expanded while midbrain marker engrail-2 (en2) was abolished by unilateral FTO depletion. Neural crest marker twist and neuronal marker n-tubulin were also decreased on the FTO morpholino injected side (Inj.). Uninjected side (Uninj.) serves as an internal control. Fractions are shown at the bottom right. Midlines of neurula embryos were marked by dashed lines. (B) HeLa cells stably expressing shRNAs targeting different sequences in FTO (#2, #3, #4; see methods) were transfected with WNT signaling reporter (TOP-flash) or control reporter (FOP-flash) (n = 3). Relative Luciferase activities (RLU) were measured after 8 hours of WNT stimulation. (C) Control HeLa cells or shFTO#4-expressing HeLa cells were stimulated with WR for 8 hours and protein levels of β-Catenin and FTO were assessed by western blot (n = 3). Note the stabilization of FTO protein by WNT signaling. (D) Control HeLa cells or shFTO#2-HeLa cells were transiently transfected with WNT reporter (TOP-flash) and co-cultured with cells expressing different shRNAs for 36 hours as indicated (n = 3). Relative luciferase activities (RLU) were measured after 16 hours of WNT stimulation. (E) Culture medium of HeLa cells transiently expressing WNT reporter (TOP-flash) was replaced with the medium harvested from control HeLa, shFTO#2-HeLa or shMETTL3-HeLa before WNT stimulation for 16 hours. RLU, Relative luciferase units. n = 3. (F) Cell lysates from Figure 2E were analysed by western blot for β-Catenin expression. GAPDH was used as a loading control for western blot. WR, WNT3A and R-Spondin1. Error bars are ± s.e.m. One-sided T-test, **: p<0.01, *: p<0.05.

To investigate the molecular mechanism how FTO regulates the WNT/β-Catenin signaling pathway, we utilized the TOP-Flash reporter harboring multiple TCF binding sites and measured the transcriptional WNT/β-Catenin signaling activity. FTO knockdown in HeLa cells inhibited WNT3A-induced activation of TOP-Flash reporter (Figure 2B), the accumulation of β-Catenin (Figure 2C), and expression of WNT/β-Catenin targets such as AXIN2 and C-MYC (Figure 2-figure supplement 1A), indicating that FTO is required for the canonical WNT/β-Catenin signaling pathway. Similar results were also found in 293T cells (Figure 2-figure supplement 1B-C) but not in HepG2 cells that express a constitutively active form of β-Catenin (Figure 2-figure supplement 1D-E), suggesting that FTO may modulate the canonical WNT signaling upstream of β-Catenin.

We then co-cultured FTO-silenced cells and TOP-Flash reporter cells to test whether FTO affects WNT/β-Catenin signaling in a cell autonomous manner. We found decreased WNT3A signaling activity even when the WNT reporter-expressing control HeLa cells were co-cultured with FTO-silenced cells (Figure 2D), demonstrating non-cell autonomous inhibition of canonical WNT signaling by FTO-depleted cells. Culture medium from FTO-depleted cell also inhibited WNT3A-induced TOP-Flash reporter activation (Figure 2E) and β-Catenin accumulation (Figure 2F), indicating that secreted factors from FTO-depleted cells antagonize the canonical WNT signaling activity. Notably, only FTO-depleted cells displayed non-cell autonomous WNT inhibition while METTL3- or YTHDF2-depleted cells showed no such activity (Figure 2D-F).

To identify the responsible factor that antagonizes the WNT/β-Catenin signaling pathway, we measured the mRNA levels of known extracellular WNT antagonists in FTO-silenced cells. Interestingly, only DKK1 mRNA was upregulated in FTO-silenced cells (Figures 3A and Figure 3-figure supplement 1A-B). Secreted DKK1 protein level was also substantially increased (Figures 3B and Figure 3-figure supplement 1C-D). When transiently depleted with DKK1 siRNA treatment, the culture medium from the FTO-depleted cells no longer inhibited the WNT3A-induced TOP-Flash reporter activation (Figure 3C), indicating that DKK1 plays a role in FTO-dependent regulation of the canonical WNT signaling pathway. Consistently, DKK1 immunodepletion from the culture medium of FTO-depleted cells also abrogated the WNT/β-Catenin inhibitory activity (Figure 3-figure supplement 1E). Similar results were found after neutralization of the DKK1 protein in the extracellular milieu by anti-DKK1 antibody (Figure 3-figure supplement 1F). These results demonstrate that secreted DKK1 proteins are responsible for the non-cell autonomous inhibition of canonical WNT/β-Catenin signaling in FTO-depleted cells.

**Figure 3.**
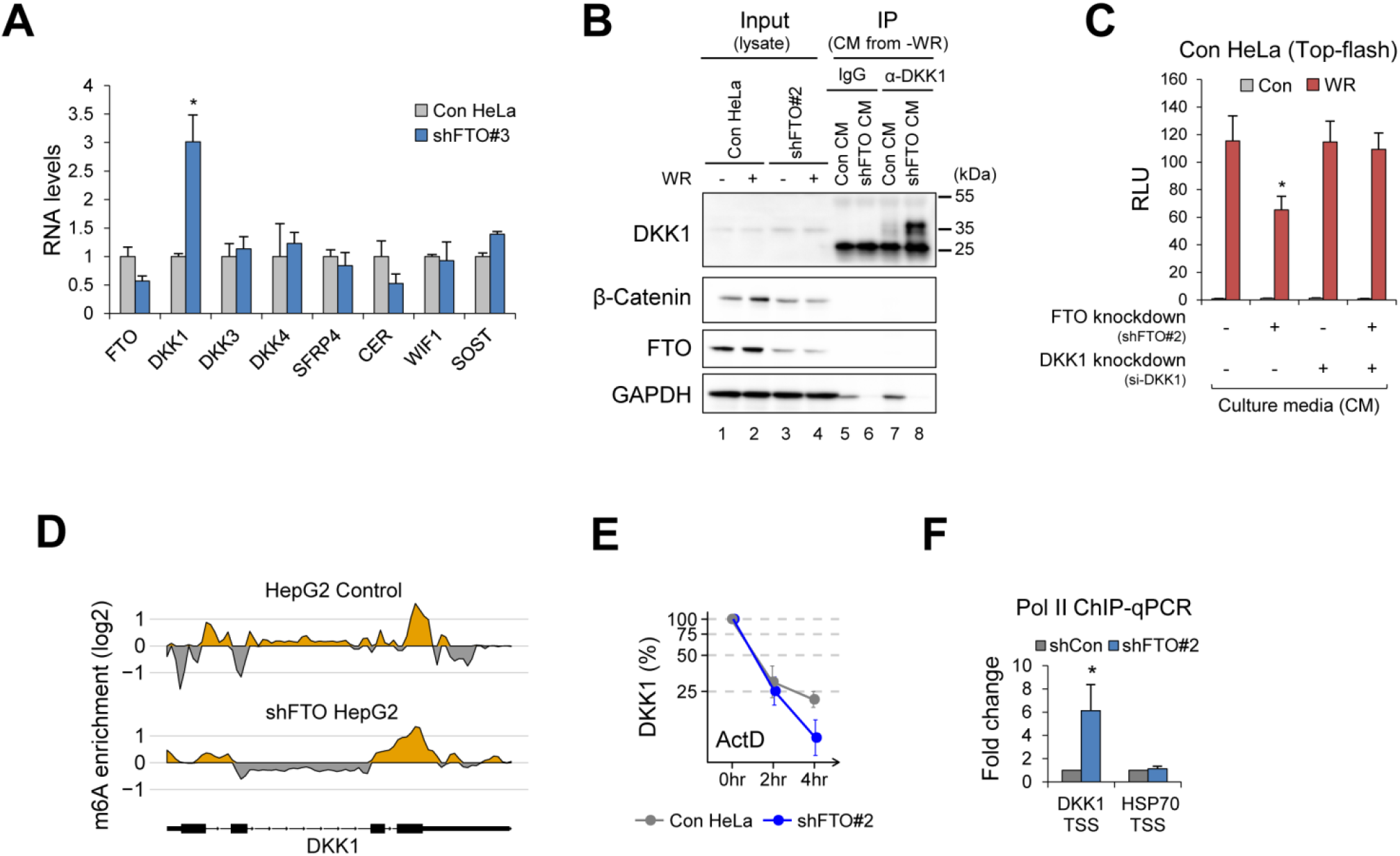
Loss of FTO increases DKK1 expression. (A) RNA levels of various extracellular WNT antagonists were measured by RT-qPCR from control or FTO-depleted HeLa cells (shFTO#3). GAPDH was used as a normalization control. n = 3. (B) Secreted DKK1 protein levels were measured by western blot after immunoprecipitating DKK1 from culture medium of control (Con CM) or FTO-depleted (shFTO#2) HeLa cells (shFTO CM). IgG immunoprecipitation serves as a negative control. (C) Culture medium of HeLa cells transiently expressing WNT reporter (TOP-flash) was replaced with the medium harvested from DKK1 siRNA or control siRNA (si-NC) transfected HeLa cells (control or FTO-depleted). Relative luciferase activities were measured after 16 hours of WNT stimulation (WR, WNT3a and R-spondin1). n = 6. (D) Change in m6A enrichment (IP/Input) across the DKK1 gene in FTO-depleted cells. Read coverages of both IP and input samples were standardized as described in Methods. The yellow areas indicate putative m6A peaks. The gene models are shown below, respectively. (E) DKK1 mRNA levels were measured by RT-qPCR from control or FTO-depleted HeLa cells treated with actinomycin D for the indicated time (n = 3). GAPDH was used as a normalization control. (F) RNA polymerase 2 (Pol II) enriched chromatin fragments were analyzed by qPCR with the primer sets encompassing DKK1 transcription start site (DKK1 TSS) or HSP90 intron 1. n = 3. GAPDH was used as a loading control for western blot. WR, WNT3A and R-Spondin1. Error bars are ± s.e.m. One-sided T-test, *: p<0.05.

FTO demethylates N6-methyladenosine (m6A) and N6, 2′-O-dimethyladenosine (m6Am) of messenger RNAs (mRNAs) and non-coding RNAs (Wei et al., 2018). m6Am has been proposed to stabilize mRNA and/or regulate cap-dependent translation (Akichika et al., 2019; Mauer et al., 2017). However, no substantial changes in m6A/m6Am modifications along the DKK1 mRNA transcript were observed in FTO-depleted HepG2 cells (Figures 3D and Figure 3-figure supplement 1G). Moreover, the stability of DKK1 mRNA were not increased after FTO knockdown (Figures 3E and Figure 3-figure supplement 1H), altogether excluding the possibility of FTO-dependent posttranscriptional regulation of DKK1 mRNA via RNA demethylation. On the contrary, Pol II occupancy near the DKK1 transcription start site (TSS) increased after FTO depletion (Figures 3F). DKK1 pre-mRNA levels were also increased in FTO-depleted cells (Figure 3-figure supplement 1I-K), indicating that the loss of FTO leads to the upregulation of DKK1 transcription. Of note, culture medium from FTO-depleted HepG2 cells, which harbor constitutively active β-Catenin, also exhibited WNT inhibitory activity in a DKK1-dependent manner (Figure 3-figure supplement 1E), excluding the possibility that the increased DKK1 expression in FTO-depleted cells is via β-Catenin-mediated transcriptional activation.

### Loss of FTO enhances cell migration via WNT/PCP pathway activation

In addition to the downregulation of canonical WNT/β-Catenin signaling in FTO-depleted cells, we also observed a consistent upregulation of the noncanonical WNT/PCP signaling across multiple cellular contexts (Figures 1B-C). WNT/PCP signaling is responsible for cell motility and polarity via Rho, Rac and JNK activation (Steffen et al., 2013). In *Xenopus*, WNT/PCP pathway regulates convergent extension (CE) movements during gastrulation, a process that elongates and shapes the body axis, in a way that either the activation or the inhibition of WNT/PCP signaling disrupts CE movements (Kim et al., 2008; Sheng et al., 2014). We found that Fto knockdown in the dorsal marginal zone produced embryos with shortened and kinked axis (Figure 4A), a hallmark of defective CE movements, suggesting possible WNT/PCP regulation by FTO.

**Figure 4.**
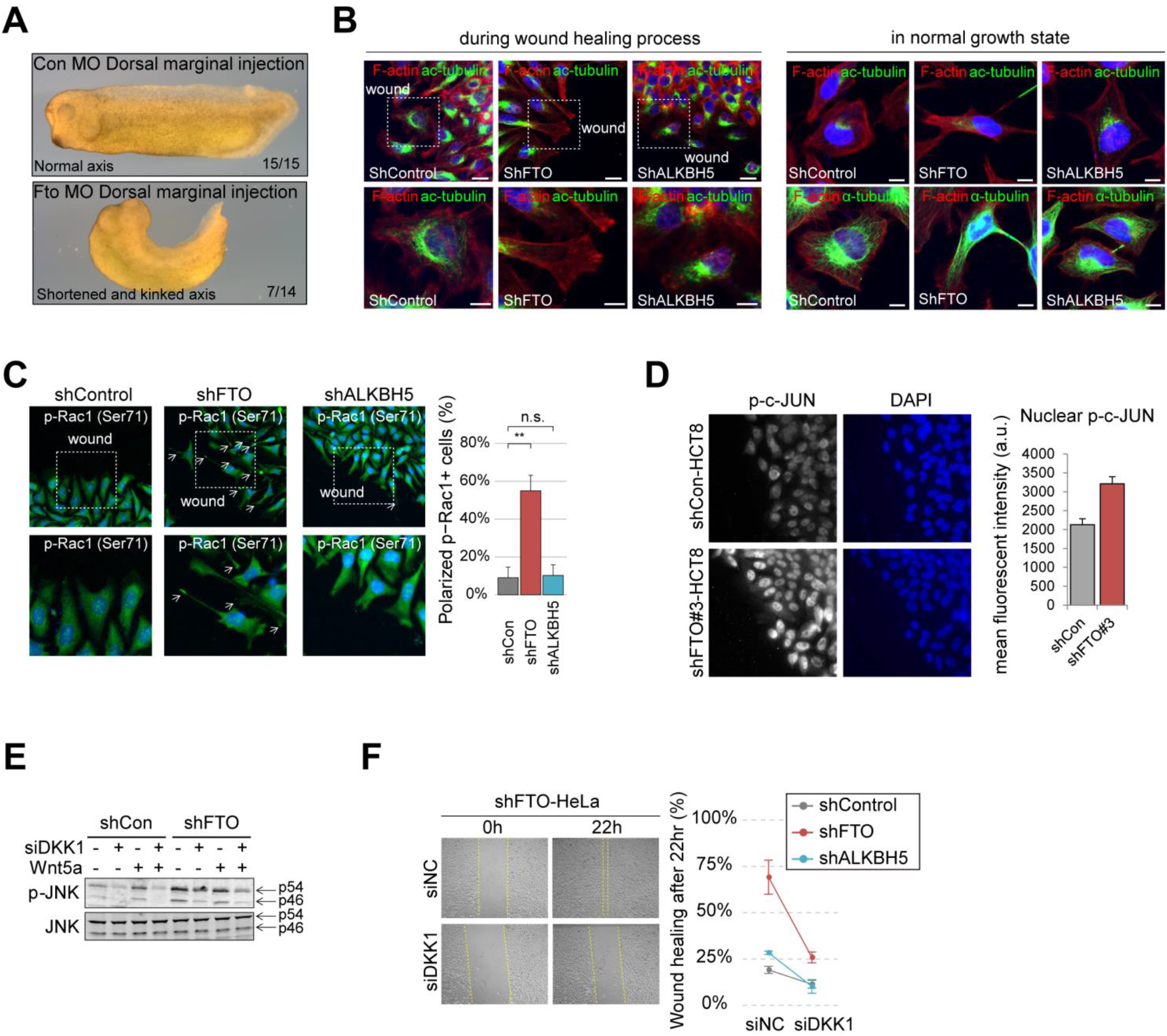
Loss of FTO enhances cell migration by activating WNT/PCP signaling. (A) Gross morphology at stage 30 tailbud embryos injected either with control morpholino (con MO) or Fto morpholino (Fto MO) at the dorsal marginal region. Fractions are shown at bottom right. Note the upper and lower images are at the same magnification. (B) Filamentous actin (F-actin) and acetylated Tubulin (ac-tubulin) were visualized in cells during wound healing (6 hours after scratch, see methods) or in normal growth state. Regions enclosed by the dashed rectangles were enlarged and shown below. Directions of wound scratch were indicated. (C) Activated Rac1 was visualized using anti-p-Rac1 (Ser71) antibody in cells after 6 hours of scratch wounding. Cells with polarized p-Rac1 staining near the wound edge were counted and graphed (right) (shCon, n = 3; shFTO, n = 4; shALKBH5, n = 3). Regions enclosed by the dashed rectangles were enlarged and shown below. Directions of wound scratch were indicated. (D) Phosphorylated c-JUN was visualized in HCT-8 cells after 6 hours of scratch wounding near the wound edge (n = 5). Mean fluorescent intensities of nuclear p-c-JUN were measured with ImageJ software and graphed (right). (E) Levels p54 and p46 JNK and its activated form (p-JNK) were assessed by western blot. Control or FTO-depleted HeLa cells were transiently transfected with either control siRNA (-) or DKK1 siRNA (+). Cells were treated with recombinant WNT5a protein for 12 hours. (F) FTO-depleted HeLa cells along with control of ALKBH5-depleted cells were further transfected with either control (siNC) or DKK1 siRNA (siDKK1). Scratch wound-healing assays were performed 48 hours after siRNA transfection. Representative wound healing of FTO-depleted cells were shown on the left. Yellow dashed lines indicate wound edges. Percentage of wound healing after 22 hours were calculated as ‘100-[wound width at 0h - wound width at 22h)/wound width at 0h]’ (n = 3). Error bars are ± s.e.m. One-sided T-test, **: p<0.01, n.s.: p>0.05.

To investigate whether and how FTO regulates WNT/PCP pathway, we first performed in vitro scratch wound healing assay to test whether there are any changes in cell motility and polarity in FTO-depleted cells. We found enhanced wound healing activity in FTO-depleted cells but not in ALKBH5-depleted cells (Figure 4-figure supplement 1A-C), indicating that FTO may play a specific role in cell migration. Moreover, FTO knockdown induced cell shape polarization at the wound edge (Figure 4B) but also in normal growth state (Figures 4B and Figure 4-figure supplement 1D), suggesting that the FTO-dependent change in cell behavior is intrinsic rather than triggered by damage response signaling during wound healing process. Of note, FTO knockdown does not seem to promote epithelial-mesenchymal transition (EMT) per se as evident by the RNA levels of EMT markers (Figure 4-figure supplement 1E). Along with the increased cell motility and polarity, active Rac1 (i.e. Ser71 phosphorylated Rac1) was accumulated at the protruding membrane of FTO-silenced cells (Figure 4C). Moreover, phophorylated c-JUN at the nucleus of FTO-silenced cells were also increased (Figure 4D), supporting the possibility of FTO-dependent WNT/PCP regulation. ALKBH5-depleted cells did not exhibit change in cell polarity and motility (Figures 4B-C, Figure 4-figure supplement 1A, Figure 4-figure supplement 1C-D).

The canonical WNT antagonist DKK1 is also known to activate the WNT/PCP signaling pathway in a number of biological contexts (Caneparo et al., 2007; Killick et al., 2014; Sellers et al., 2018). Thus, we hypothesized that the FTO-dependent upregulation of DKK1 transcription (Figures 3 and Figure 3-figure supplement 1) may also be responsible for WNT/PCP activation. To test this hypothesis, we treated FTO-depleted cells with WNT5A, a known WNT/PCP ligand (Ohkawara et al., 2011). Indeed, FTO knockdown augmented phosphorylation of JNK and c-JUN (Figures 4D-E and Figure 4-figure supplement 1F), which is the hallmark of WNT/PCP activation (Yamanaka et al., 2002). This increased JNK phosphorylation was abrogated after DKK1 knockdown using siRNA (Figure 4E), indicating that the upregulation of DKK1 is responsible for the WNT/PCP activation in FTO-depleted cells. DKK1 knockdown also suppressed the enhanced cell migration of FTO-depleted cells (Figure 4F), demonstrating the role of FTO-dependent DKK1 upregulation in cell migration via WNT/PCP activation.

### WNT/STOP pathway induces FTO protein accumulation in the cytosol to regulate the stability of m6A mRNA targets

Unexpectedly, we found that WNT3A stimulation leads to the accumulation of FTO proteins (Figure 2C), suggesting that the canonical WNT pathway may also regulate the function of FTO. Canonical WNT signaling is known to stabilize proteins other than β-Catenin by inhibiting GSK3 kinase (Acebron et al., 2014; Kim et al., 2015; Taelman et al., 2010), collectively known as the WNT/STOP (<u>ST</u>abilization <u>O</u>f <u>P</u>roteins) pathway (Acebron et al., 2014). While FTO was not reported as a target of the WNT/STOP pathway, it was recently reported that GSK3 can phosphorylate FTO and generate phosphodegron for polyubiquitination and subsequent proteasomal degradation (Faulds et al., 2018). Consistent with previous reports, both WNT3A stimulation and GSK3 inhibition (i.e. SB216763) induce FTO protein stabilization by inhibiting polyubiquitination (Figures 5A and Figure 5-figure supplement 1A), indicating that the canonical WNT/STOP signaling also stabilizes the FTO protein via GSK3 inhibition.

**Figure 5.**
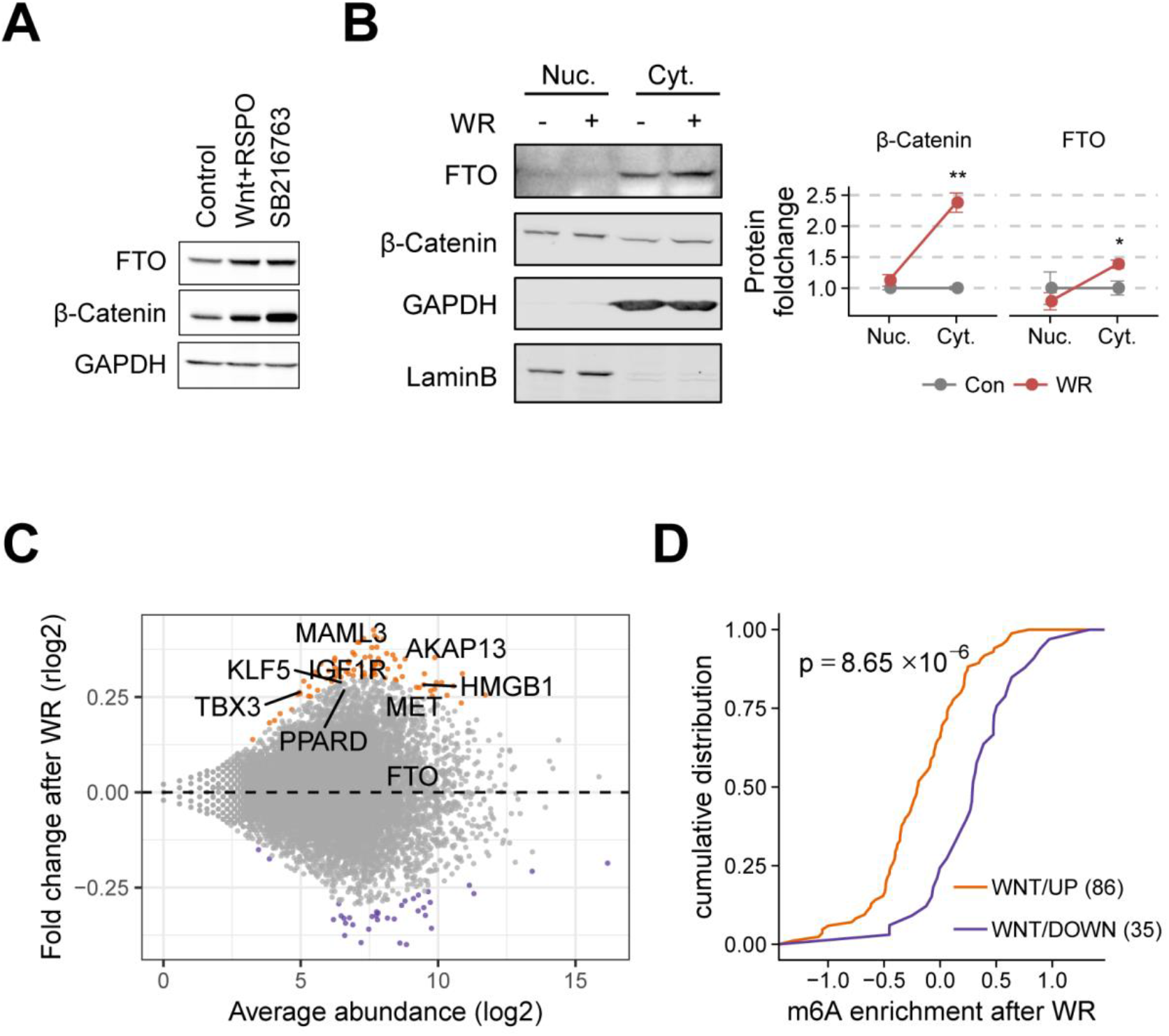
Canonical WNT induces mRNA stabilization via m6A demethylation. (A) FTO protein levels were measured in WNT stimulated (WNT+RSPO, WNT3a and R-spondin1) or GSK3 inhibited (SB216763) HeLa cells. β-Catenin levels were used as a control for WNT activation or GSK3 inhibition. GAPDH was used as a loanding control. (B) Cytosolic FTO and β-Catenin protein levels after WNT stimulation (WR, WNT3A and R-spondin1) were assessed along with nuclear fractions (nuclear fraction, n = 3; cytosolic fraction, n = 6). Protein levels were quantified and graphed (right). GAPDH-normalized cytosolic fractions and LaminB-normalized nuclear fractions were calculated. P values are indicated. (C) MA plot of regularized log2 fold change (rlog2) and average abundance after WNT3a treatment (WR). Up-regulated genes are in orange, and down-regulated genes are in purple (p-value = 0.15). Canonical WNT targets that were differentially expressed are labelled (p-value = 0.25). (D) Cumulative plot of m6A enrichment of up-regulated genes (orange) and down-regulated genes (purple) after WNT3a treatment. The number of genes are shown in parenthesis. One-sided Kolmogorov-Smirnov test was used to test statistical significance.

The subcellular distribution of FTO proteins determines the substrate repertoire of RNA demethylation (Wei et al., 2018). Therefore, we measured the WNT3A-induced FTO accumulation in different subcellular fractions to investigate the functional implication of FTO protein stabilization. Interestingly, FTO proteins were significantly accumulated in the cytosolic fraction but not in the nuclear fraction (Figure 5B). This result hints at the possible canonical WNT/STOP mediation of internal m6A and cap m6Am of poly(A) RNAs in the cytosol. To test whether canonical WNT signaling mediates m6A(m) methylation, we performed m6A-seq in HeLa cells after WNT3A stimulation. 86 genes were up-regulated including known canonical WNT targets such as MET and TBX3 (p-value < 0.15, Figure 5C). When comparing the m6A enrichment of up- and down-regulated genes after WNT3A stimulation, we found that upregulated genes were less methylated than downregulated genes (p-value=8.65 × 10^−6^, Figure 5D), suggesting that canonical WNT/STOP signaling may regulate gene expression via RNA demethylation. Of note, WNT3A stimulation and GSK3 inhibition did not stabilize other m6A pathway components such as the m6A writer METTL3 (Liu et al., 2014), m6A reader YTHDF2 (Wang et al., 2014) and ALKBH5 (Zheng et al., 2013) (Figure 5-figure supplement 1B). mRNA levels of m6A pathway components including FTO were also unaffected after WNT3A stimulation or GSK3 inhibition (Figure 5-figure supplement 1C).

Internal m6A methylation is known to reduce the stability of poly(A) mRNAs (Wang et al., 2014), and so FTO-dependent demethylation promotes mRNA stability (Wei et al., 2018). In fact, known WNT/β-Catenin target genes were overall downregulated after FTO knockdown in HepG2 cells, where β-Catenin is constitutively activated (Figure 6-figure supplement 1A), hinting at the possibility of FTO-dependent posttranscriptional regulation of WNT/β-Catenin target genes. To select candidates for WNT-dependent posttranscriptional regulation, we manually examined the standardized m6A read coverage of upregulated genes after WNT3A stimulation. Specifically, known canonical WNT/β-Catenin target MET, but not TBX3, exhibited a substantial reduction in m6A modification upon WNT3A stimulation (Figure 6A and Figure 6-figure supplement 1B). We also found a similar reduction in m6A modification for HMGB1, AKAP13, MAML3 and IGF1R (Figures 6B-C and Figure 6-figure supplement 1C-D). mRNAs of these posttranscriptional WNT targets, including MET, were more stabilized by WNT3A in a FTO-dependent manner (Figures 6A-C and Figure 6-figure supplement 1E-F). AXIN2, a well-known transcriptional WNT target, was also posttranscriptionally stabilized by WNT3A in a FTO-dependent manner (Figure 6-figure supplement 1G). Overexpression of FTO also induced mRNA stabilization of these posttranscriptional WNT targets (Figure 6D), supporting the FTO dependency. Taken together, these results indicate that canonical WNT/STOP signaling stabilizes FTO proteins in the cytosol to regulate the stability of mRNA via m6A demethylation.

**Figure 6.**
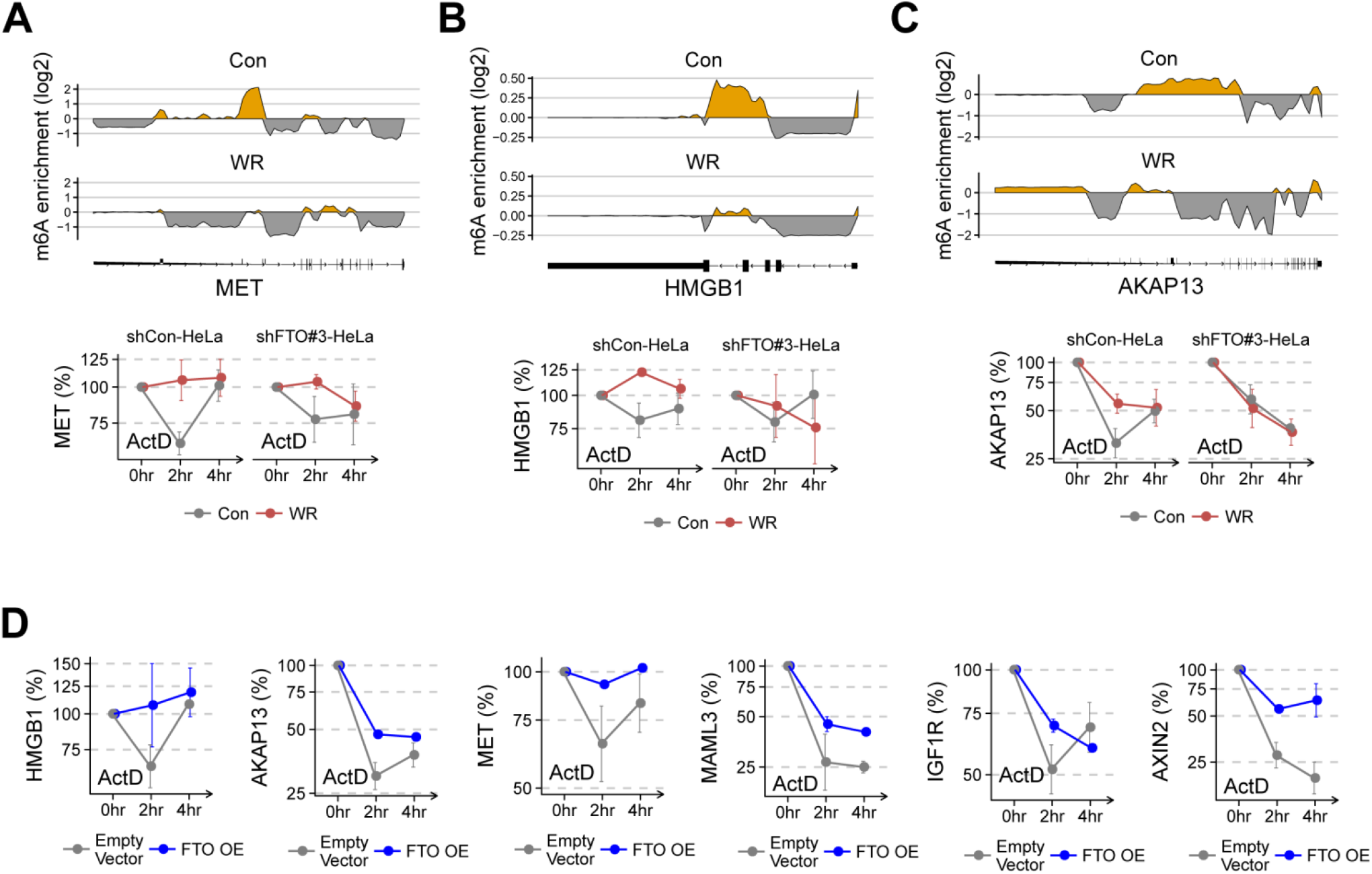
Canonical WNT stabilizes FTO target mRNAs. (A-C) Change in m6A enrichment (IP/Input) and mRNA decay curves across the MET (A), HMGB1 (B) and AKAP13 (C) genes after WNT3A treatment (WR). Read coverages of both IP and input samples were standardized as described in Methods. The yellow areas indicate putative m6A peaks. The gene model for each genes are shown below the coverage plot. mRNA levels of each genes were measured by RT-qPCR in control of FTO-depleted HeLa cells treated with actinomycin D (act D) for the indicated time. Mock treated controls were marked with dashed lines and WNT treated samples were shown with solid lines (n = 3). (D) RNA levels of HMGB1, AKAP13, MET, MAML3, IGF1R and AXIN2 were measured by RT-qPCR in empty vector or FTO (FTO OE) tranfected HeLa cells treated with actinomycin D (act D) for the indicated time (n = 2). GAPDH was used as a normalization control. Error bars are ± s.e.m.

## DISCUSSION

In this study, we applied a functional genomics approach to characterize FTO-depleted cells in a data-driven manner. In particular, we identified multiple functional links between FTO and WNT signaling pathways. We found that loss of FTO non-cell autonomously inhibits the canonical WNT/β-Catenin pathway but also enhances the non-canonical WNT/PCP pathway via transcriptional upregulation of DKK1 (Figure 6-figure supplement 1H). Surprisingly, cytoplasmic FTO proteins were accumulated upon canonical WNT signaling in a GSK3-dependent manner (Figure 5A-B). m6A writer METTL3 and the other m6A erasers ALKBH5 were unaffected (Figure 5-figure supplement 1B-C), suggesting that this accumulation of FTO is responsible for the epitranscriptome change upon canonical WNT signaling (Figures 5C-D). Specifically, we identified the m6A demethylation and subsequent stabilization of the mRNA of MET, HMGB1, AKAP13, MAML3 and IGF1R upon canonical WNT signaling (Figures 6A-C, Figure 6-figure supplement 1C-F). It is worth mentioning that FTO is also associated with the noncanonical WNT/Ca+ pathway (Osborn et al., 2014). While the exact molecular factor responsible for this association remains unknown, the report supports our observation of this intricate entanglement between FTO and multiple WNT signaling pathways.

WNT response is well established in the context of transcriptional (e.g. WNT/β-Catenin and WNT/Ca2+) and posttranslational (e.g. WNT/STOP and WNT/PCP) regulation (Acebron et al., 2014; Katoh, 2017; Lai et al., 2009; MacDonald et al., 2009). However, the direct impact of WNT signaling in RNA regulation has not been widely appreciated and mostly considered anecdotal. For instance, upon canonical WNT signaling, cytoplasmic HuR proteins recognize the AU-rich element of the 3′ UTR of Pitx2 mRNA and enhance the stability of Pitx2 mRNA to regulate cell-type-specific proliferation during mammalian development (Briata et al., 2003). Here, we characterize the molecular response to canonical WNT signaling in the context of RNA modifications. Specifically, we identify the FTO protein as another target of the WNT/STOP pathway and discover the subsequent FTO-dependent mRNA stabilization by m6A demethylation. Indeed, the physiological meaning of WNT-dependent RNA methylation warrants further investigation, particularly in the context of cytoplasmic accumulation of FTO proteins.

The multifacetedness of WNT signaling is often attributed to the opposing effects of WNT proteins (e.g. WNT3a and WNT5a), WNT receptors (e.g. FZD family of G protein couple receptors), co-receptors (e.g. LRP6 and ROR2), and its dual regulators of the canonical and non-canonical pathways. For example, Dishevelled (DVL) proteins interact with GSK3 to inhibit β-Catenin protein degradation, but also play a role in the activation of JNK, AP-1, and other components of the noncanonical WNT pathway (Boutros et al., 1998; Habas et al., 2003; Li et al., 1999). Frat oncoproteins are reported to behave in a similar manner as DVL (van Amerongen et al., 2010). We extend this list of branchpoint regulators by investigating multiple WNT signaling pathways in FTO-depleted cells. In particular, we find that FTO regulates the expression of DKK1, another dual WNT regulator (Grumolato et al., 2010; Sellers et al., 2018), thereby contributing to the bifurcation of the canonical and noncanonical pathways. While the exact molecular mechanism of FTO-dependent DKK1 regulation remains not completely understood, we find that it ultimately affects DKK1 transcription. In all, this study proposes DKK1-mediated WNT signaling as a plausible explanation underlying the various functions of FTO in development and human diseases.

## MATERIALS and METHODS

### Plasmids, oligonucleotides, antibodies and reagents

Wildtype Flag-hFTO and its catalytic mutant Flag-hFTO (HD) were gifts from Dr. Chuan He. pRK5-HA-ubiquitin (Addgene #17608) was a gift from Dr. T. M. Dawson (Lim et al., 2005). Control short hairpin RNA (shRNA) in pLKO.puro backbone was a gift from Dr. David Sabatini (Addgene #1864). ShRNA oligonucleotides against human FTO, METTL3, YTHDF2 and ALKBH5 were synthesized (Cosmogenetech) and ligated to pLKO1.puro (Addgene #8453). Sequences of shRNA oligonucleotides are as follows:

shFTO#2 F:

5’-CCGGCGGTTCACAACCTCGGTTTAGCTCGAGCTAAACCGAGGTTGTGAACCGTTTTTG-3’

shFTO#2 R:

5’-AATTCAAAAACGGTTCACAACCTCGGTTTAGCTCGAGCTAAACCGAGGTTGTGAACCG-3’

shFTO#3 F

5’-CCGGCCCATTAGGTGCCCATATTTACTCGAGTAAATATGGGCACCTAATGGGTTTTTG-3’

shFTO#3 R

5’-AATTCAAAAACCCATTAGGTGCCCATATTTACTCGAGTAAATATGGGCACCTAATGGG-3’

shFTO#4 F

5’-CCGGACCTGAACACCAGGCTCTTTACTCGAGTAAAGAGCCTGGTGTTCAGGTTTTTTG-3’

shFTO#4 R

5’-AATTCAAAAAACCTGAACACCAGGCTCTTTACTCGAGTAAAGAGCCTGGTGTTCAGGT-3’

shALKBH5 F:

5’-CCGGCCTATGAGTCCTCGGAAGATTCTCGAGAATCTTCCGAGGACTCATAGGTTTTTTG-3’

shALKBH5 R:

5’-AATTCAAAAACCTATGAGTCCTCGGAAGATTCTCGAGAATCTTCCGAGGACTCATAGG-3’

shMETTL3#2 F

5’-CCGGGCCAAGGAACAATCCATTGTTCTCGAGAACAATGGATTGTTCCTTGGCTTTTTG-3’

shMETTL3#2 R

5’-AATTCAAAAAGCCAAGGAACAATCCATTGTTCTCGAGAACAATGGATTGTTCCTTGGC-3’

shYTHDF2#1 F

5’-CCGGGCTACTCTGAGGACGATATTCCTCGAGGAATATCGTCCTCAGAGTAGCTTTTTG-3’

shYTHDF2#1 R

5’-AATTCAAAAAGCTACTCTGAGGACGATATTCCTCGAGGAATATCGTCCTCAGAGTAGC-3’

Small interfering RNAs (siRNAs) duplexes were synthesized (Bioneer) and used. Sequences of siRNAs are as follows:

si-DKK1: 5’-CCUGUCCUGAAAGAAGGUCAA-3’ (sense strand)

si-control: 5’-CCUACGCCACCAAUUUCGU-3’ (sense strand)

Primary antibodies used were anti-β-Catenin (Santa Cruz #sc-7963), anti-acethylated Tubulin (Sigma #T7451), anti-alpha Tubulin (Abcam #ab52866), anti-FTO (Abcam #ab124892), anti-GAPDH (Santa Cruz #sc-32233 and #sc-25778), anti-m6A (SYSY #202003), anti-RNA Polymerase 2 (Santa Cruz #sc-56767), anti-DKK1 (R&D systems #AF1096 and Santa Cruz #sc-374574), anti-p-JNK (Cell Signaling #4668), anti-JNK (Cell Signaling #9258), anti-p-Rac1 (Ser71) (Cell Signaling #2461), anti-LaminB (Abcam #ab16048), anti-Axin2 (Cell Signaling #2151), anti-C-Myc (Cell Signaling #5605), anti-p-c-Jun (Santa Cruz #sc-822), anti-METTL3 (Abcam #ab195352), anti-YTHDF2 (Abcam #ab170118) and anti-ALKBH5 (Abcam #ab69325).

Recombinant murine Wnt3a (PeproTech #315-20), human/mouse Wnt5a (R&D Systems #645-WN-010) and human R-Spondin1 (R&D Systems #4645-RS-025) were purchased and used. GSK3 inhibitor SB216763 (Sigma #S3442) was dissolved in DMSO and used at 10 μM. Alexa Fluor^™^ 647 Phalloidin (Invitrogen #A22287) was used to stain F-actin.

### Cell culture, Lentiviral shRNA delivery and transfection

HeLa, 293T, HepG2 and SW480 cells were cultured in DMEM containing 10% FBS. Lentiviral supernatants were produced using Lenti-X Packaging Single Shots (VSV-G) (Clontech #631275) mixed with pLKO.1 puro-based lentiviral constructs transfected to Lenti-X 293T cells. 24 hours after transfection, supernatants were collected and concentrated using Lenti-X Concentrator (Clontech #631231). Stable shRNA-expressing cell lines were established by infecting cells with shRNA-containing lentiviral supernatants together with 4 μg/ml protamine sulfate for 24 hours. After 24-48 hours of culture with fresh medium, cells were selected with medium containing 1∼2 μg/ml puromycin (AG Scientific #P-1033) until non-transduced cells became completely dead, which usually takes 2-3 days. Plasmid DNAs were transfected using Fugene HD (Promega #E2312) according to the manufacturer’s protocol.

### WNT reporter assays

Luciferase activities were measured using Dual-Luciferase assay system (Promega #E1980) according to the manufacturer’s protocol. WNT reporter plasmid (TOP-flash) or control reporter (FOP-flash) together with CMV-Renilla plasmid were transiently transfected to shRNA-expressing cells. For co-culture experiments, reporter plasmids were transfected to either control or shFTO-expressing HeLa cells. After 24 hours of transfection, transfected cells were co-cultured with shRNA-expressing cells with 1:1 ratio. Mixed cells were harvested after additional 24 hours and luciferase activities were measured. For CM (culture medium or conditioned medium) experiments, culture media were harvested from shRNA-expressing cell cultures at least after 24 hours of media change. Collected media were then filtered using syringe filter (Millipore #SLGP033RS) before applying to the WNT reporter-transfected cell cultures. For immunodepletion of DKK1 protein, collected and filtered culture medium was incubated with anti-DKK1 antibody or with normal IgG overnight at 4 °C. Antibody-protein complex was then removed with excessive protein A/G agarose bead.

### *Xenopus* embryos manipulation

*Xenopus laevis* were obtained from the Korean *Xenopus* Resource Center for Research and from Nasco (Wisconsin, USA). Embryo collection, microinjection and in situ hybridization were performed as described (Kim et al., 2009). Developmental stages were determined as previously described (Nieuwkoop, 1967). *Xenopus* maintenance and embryo experiments were carried out with protocols approved by Seoul National University Institutional Animal Care and Use Committees (Seoul, Korea).

### RNA isolation, quantitative RTPCR

Total RNAs were extracted with TRIzol (Invitrogen) and treated with DNase I (Takara). 200-500 ng of total RNAs were reverse transcribed using Primescript RT master mix (Takara). Quantitative PCR was performed using Power SYBR Green PCR Master Mix (Thermo Fisher Scientific) under StepOnePlus Real-Time PCR system (Thermo Fisher Scientific). Primer sequences are listed in Table 1.

**Table 1.**
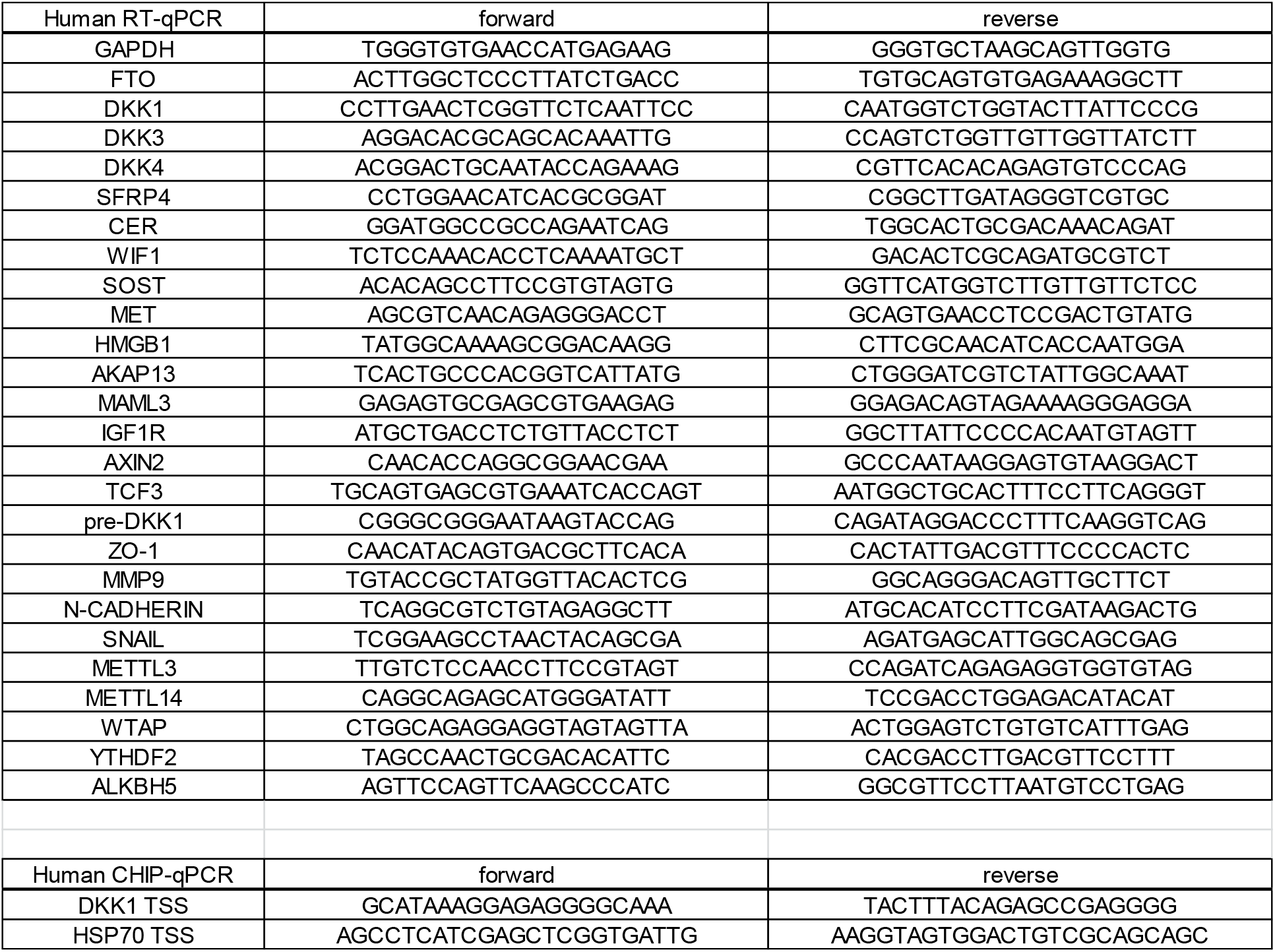
qPCR primers used in the study.

### Western blot and ubiquitination assay

For western blots, cells were lysed with a lysis buffer containing 20 mM HEPES, pH 7.2, 150 mM NaCl, 1% Triton X-100, 10 mM EDTA, pH 8.0, 1 mM DTT, together with protease inhibitor cocktail (Promega) and phosphatase inhibitor cocktail (Promega). Cell lysates were separated on Novex WedgeWell Tris-Glycine Mini Gels (Thermo Fisher Scientific) and transferred to polyvinylidene fluoride membrane (GE Healthcare) or to nitrocellulose membrane (Amersham). Blots were washed with PBST (PBS containing 0.1% Tween-20), blocked with 3% BSA for 1 hour at room temperature and incubated with primary antibodies overnight at 4 °C. Membranes were then washed three times with PBST and incubated with secondary antibodies for 1 hour at room temperature. Secondary antibodies were from Li-Cor (IRDye 680 and IRDye 800, 1:10,000) for infrared imaging of proteins using the LiCor Odyssey system. For chemiluminiscence detection of signals, HRP-conjugated secondary antibodies (Jackson Laboratories) were used.

Ubiquitination assays were performed as described (Kim et al., 2015). Briefly, HeLa cells were transfected with Flag-FTO together with HA-ubiquitin. After 48 hours of transfection, cells were treated either with DMSO or with SB-216763, together with a proteasomal inhibitor MG132 for 2 to 4 hours. Cells were then lysed with lysis buffer containing 20 mM HEPES, pH 7.2, 150 mM NaCl, 1% Triton X-100, 10 mM EDTA, pH 8.0, 1mM DTT and 1 mM NaF together with a protease inhibitor cocktail and incubated in Eppendorf tubes at 95°C in the presence of 1% SDS to disrupt protein-protein interactions. The lysates were then diluted 10-fold to reduce SDS concentration to 0.1% before immunoprecipitating with Flag antibody conjugated agarose beads (Sigma #A2220). Bead-lysate mixtures were incubated overnight at 4°C using an end-over-end rotator. Beads were then washed three times with cold buffer (10 mM Tris, pH 9.0, 100 mM NaCl, 0.5% NP40) followed by elution with SDS loading buffer at 95°C for 5 minutes.

### Methylated RNA immunoprecipitation and m6A-seq library preparation

Polyadenylated RNAs (polyA+ RNAs) were prepared using Dynabeads mRNA DIRECT kit (ambion). PolyA+ RNAs were then fragmented using fragmentation buffer (Ambion #AM8740) to yield ∼200 nt lengths RNA fragments. Polyclonal anti-m6A antibody (Synaptic Systems) was incubated with fragmented RNAs in RIP buffer (10 mM Tris-HCl pH 7.5, 150 mM NaCl, 0.1% NP40) supplemented with RNase inhibitors, RNasin (promega) and Superase-in (Ambion), for 4 hours at 4 °C. Along with this, Dynabeads protein A (Invitrogen #10002D) was washed and pre-blocked with 0.5 mg/ml BSA for 2 hours at 4 °C. Beads were then added to antibody-RNA complex and incubated for an additional 1 hour at 4 °C. Bound RNA was eluted after extensive washing with RIP buffer by competitive elution with buffer containing 7 mM m6A (Abcam #ab145715) twice each for 1 hour at 4 °C. Eluted RNAs were then precipitated with ethanol and used for m6A-sequencing library preparation.

Sequencing libraries were prepared using SMARTer Stranded RNA-seq kit (Clontech #634837) according to the manufacturer’s protocol. Purified m6A-seq libraries were sequenced on an Illumina MiSeq platform with 10-20% PhiX control library (Illumina). The data is deposited in the Gene Expression Omnibus (GEO) under the accession numbers GSEXXXXXX.

### Chromatin immunoprecipitation

For chromatin immunoprecipitation with RNA polymerase 2 antibody or with FTO antibody, cells were first fixed and crosslinked with 1% formaldehyde for 10 minutes and quenched with 0.25 M glycine before lysed with RIPA lysis buffer (50 mM Tris-HCl pH 7.5, 150 mM KCl, 5 mM EDTA, 1% NP40, 0.1% SDS and 0.5% sodium deoxycholate supplemented with cocktails of protease inhibitor and phosphatase inhibitor) and sonicated to obtain DNA fragments around 500 bp. These samples were then precleared with pre-saturated protein A bead with yeast tRNA and salmon sperm DNA followed by the incubation with antibody-bead mixture overnight at 4 °C. After extensive washing with RIPA wash buffer (50 mM Tris-HCl pH 7.5, 150 mM KCl, 1% NP40, 0.25% sodium deoxycholate) and TE buffer (10 mM Tris-HCl pH 7.5, 1 mM EDTA), samples were resuspended in RIPA elution buffer (50 mM Tris-HCl pH 7.5, 5 mM EDTA, 10 mM DTT, 1% SDS) and reverse crosslinked overnight at 65 °C. DNA fragments were purified using Qiaquick PCR purification kit (Qiagen) after proteinase K digestion for 1 hour at 45 °C. RNA polymerase 2 or FTO bound DNA fragments were quantified by quantitative PCR using primer sets listed in Table 1.

### Scratch wound healing assay

For scratch wound healing assay, cells were grown near confluency before making scratch wound using 1-200 ul micropipette tips. For positional reference, bottom side of culture plate was marked. Wound healing acvitities were traced and photographed under stereomicroscope. For immunocytochemistry near wound edge, cells were grown on the coverslip near confluency and scratch wound was introduced as described above.

### Immunocytochemistry

For immunocytochemistry, cells were grown on top of coverslip and fixed with 4% formaldehyde for 1 hour at room temperature. Fixed cells were washed and permeabilized with PBST and blocked with 3% BSA for 1 hour at room temperature. Primary antibodies were applied overnight at 4°C. After 1 hour of secondary antibody incubation and extensive PBS washing, coverslips were mounted on the slide glass with Prolong Gold antifade reagent with DAPI (Life Technologies #P-36931) and visualized under Zeiss LSM 700 confocal microscope.

### Sequence data analysis

Pre-processed siFTO and siALKBH5 knockdown RNA-seq data were downloaded from GEO:GSE78040 (Mauer et al., 2017). Pre-process shFTO knockdown RNA-seq data on HepG2 and K562 cells were downloaded from the ENCODE portal (Consortium, 2012; Davis et al., 2018) (https://www.encodeproject.org/) the following identifiers: ENCFF019JYU, ENCFF375TPO, ENCFF605EOA, ENCFF622ZUI, ENCFF041CFW, ENCFF739AIP, ENCFF826FQP, and ENCFF930LYD. BaseMean, regularized fold-change, and differential gene expression analysis were conducted using DESeq2 v1.14.1 (Love et al., 2014) for genes with more than 1 aligned read. Transcriptional targets of LEF1, TCF7, ATF2, and ATF4 were from TFactS (Essaghir et al., 2010) and download on March 6th 2019. Canonical WNT targets were downloaded from the WNT homepage (https://web.stanford.edu/group/nusselab/cgi-bin/wnt/target_genes) and downloaded on Nov 21st 2018.

### Gene set enrichment analysis

Gene set enrichment scores were computed using the PAGE algorithm (Kim and Volsky, 2005). Briefly, the Central Limit Theorem states that the distribution of the average of randomly sampled n observations approaches the normal distribution for larger n. PAGE utilizes this theorem to model the null distribution and estimate the effect size (i.e. Z score) and p-value of a given set of genes. Specifically, given the regularized fold change of gene g_i_ and a gene set S = (g_1_, g_2_, …, g_n_), the gene set enrichment score z_s_ is the following:

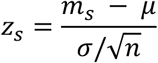

Where m_s_ is the sample mean, μ is the population mean, and σ is the population variance. 487 Wikipathway genesets (Kutmon et al., 2016) were downloaded on March 7th 2019. Mouse Gene Ontology terms and gene sets (Ashburner et al., 2000; Carbon et al., 2009; The Gene Ontology, 2019) were downloaded on April 5th 2016. Only gene sets of at least size 10 were used for further analysis.

### m6A-seq data processing and analysis

m6A-seq reads were mapped to the human GRCh38 genome using STAR v2.5.2b with default options (Dobin et al., 2013). Read coverage of aligned reads was computed using samtools 1.8 (Li et al., 2009) and bedtools 2.26 (Quinlan and Hall, 2010). Custom standardization of m6A-seq read coverages were applied as done previously (Kim et al., 2021). Briefly, to account for the technical variation in sample and library preparation, input and IP read coverages for each gene were first normalized by the pseudo-median (Hodges, 1963) and then smoothened by natural cubic splines with degree of freedom = 50. Natural cubic splines were computed using the splines R package.

## ACKNOWLEDGEMENTS

We thank Chuan He for FTO plasmids; Jaechul Lim for help in m6A-seq library preparation; Jeongwoon Lee for assistance in cell migration assays; Da-Eun Choi, Eunji Kim and Jihye Yang for technical support and help in plasmids construction; Jae Ho Paek, Jong Woo Bae, Yongwoo Na, Sha Yu, Yoosik Kim, Wantae Kim and Eek-hoon Jho for critical reading of the manuscript; Narry Kim for discussion and general supervision. We thank the ENCODE Consortium and the Brenton Graveley lab for generating the shFTO RNA-seq datasets. We also thank members of Narry Kim’s laboratory for discussions and comments. This work was supported by IBS-R008-D1 of the Institute for Basic Science from the Ministry of Science and ICT of Republic of Korea.

## FIGURE SUPPLEMENTS

**FIGURE 1-figure supplement 1.**
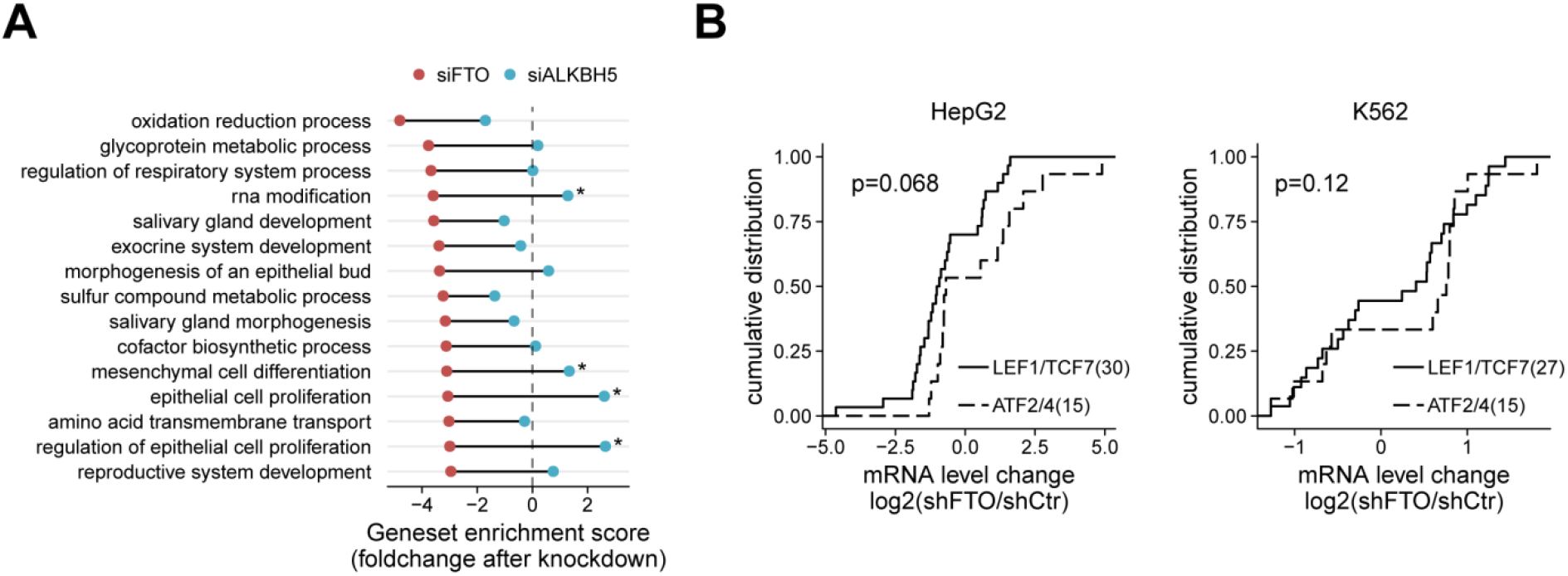
Functional genomics analysis reveals roles of FTO in the regulation of multiple WNT pathways. (A) Top 15 mouse Gene Ontology biological process gene sets with greater negative enrichment score in siFTO-depleted cells (red dots) than in siALKBH5-depleted cells (blue dots). Genesets with enrichment score difference greater than 4 are marked. (B) Cumulative plot of LEF1/TCF7 targets (solid) and ATF2/ATF4 targets (dashed) after FTO knockdown in HepG2 (left) and K562 (right) cells. Only differentially expressed targets (p-value < 0.01 and p-value < 0.05, respectively) were chosen. Of note, P-value thresholds were chosen to achieve comparable gene set size. The number of genes are shown in parenthesis. One-sided Kolmogorov-Smirnov test was used to test statistical significance.

**FIGURE 2-figure supplement 1.**
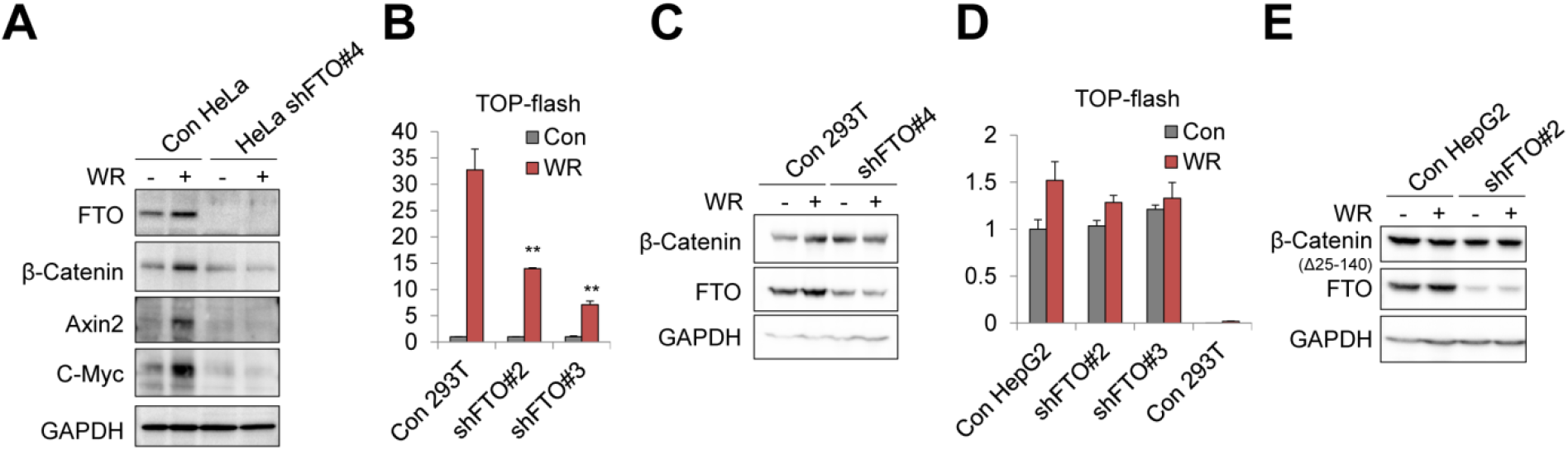
FTO is required for the canonical WNT/β-Catenin signaling. (A) Protein levels of FTO, β-Catenin, Axin2 and C-Myc were assessed in control or FTO-depleted HeLa cells treated with WNT3a and R-spondin1 (WR) by western blot. GAPDH was used as a loading control. (B) Control or FTO-depleted (shFTO#2 and shFTO#3, see methods for details) 293T cells transiently expressing WNT reporter (TOP-flash) were stimulated with WNT (WR, WNT3a and R-spondin1) for 16 hours. Relative luciferase activities were measured. n = 3. (C) Control or FTO-depleted (shFTO#4) 293T cells were stimulated with WNT (WR, WNT3a and R-spondin1) for 16 hours. Protein levels of β-Catenin and FTO were measured by western blot. GAPDH was used as a loading control. (D) Control or FTO-depleted (shFTO#2 and shFTO#3) HepG2 cells transiently expressing WNT reporter (TOP-flash) were stimulated with WNT (WR, WNT3a and R-spondin) for 16 hours. Relative luciferase activities were measured and compared with the results in control 293T cells. n = 3. (E) Control or FTO-depleted HepG2 cells were stimulated with WNT signaling and protein levels of β-Catenin and FTO were measured by western blot. GAPDH was used as a loading control. Truncated form of β-Catenin (∼70 kDa, Δ25-140) was detected in HepG2 cells as previously described. Error bars are ± s.e.m. One-sided t-test. **: p<0.01.

**FIGURE 3-figure supplement 1.**
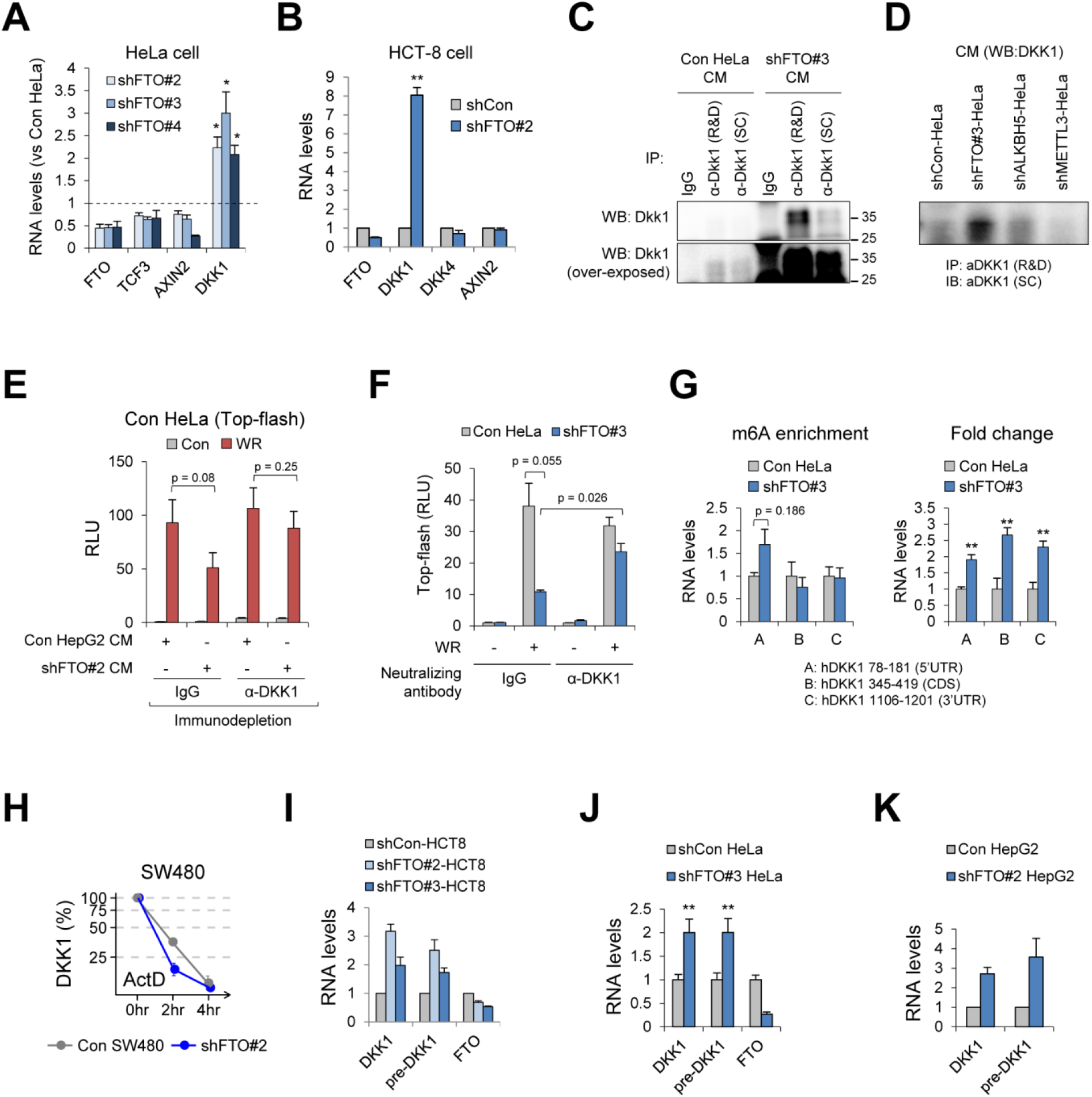
FTO regulates DKK1 RNA at the transcriptional level. (A) RNA levels of FTO, TCF3, AXIN2 and DKK1 were measured in FTO-depleted HeLa cells (shFTO#2, shFTO#3 and shFTO#4, see methods for details) and normalized with control HeLa cells (shFTO#2, n = 5; shFTO#3, n = 3; shFTO#4, n = 4). GAPDH was used for the RT-qPCR normalization. (B) RNA levels of FTO, DKK1, DKK4 and AXIN2 in HCT-8 cells stably expressing either shCon or shFTO#2 (see methods for details). GAPDH was used for the RT-qPCR normalization. n = 3. (C) Two different DKK1 antibodies were compared for the detection of secreted DKK1 protein in the culture medium from control or FTO-depleted HeLa cells. DKK1 antibody from R&D systems (R&D) resulted better immunoprecipitation than the DKK1 antibody from Santa Cruz Biotech. (SC). DKK1 (SC) was used for western blot. IgG was used as a negative control. (D) Secreted DKK1 protein levels were assessed by immunoprecipitation of DKK1 using anti-DKK1 antibody (R&D) from the culture media of control, FTO-, ALKBH5- or METTL3-depleted HeLa cells, followed by western blot with anti-DKK1 antibody from Santa Cruz (SC). (E) Culture medium of WNT reporter expressing HeLa cells were replaced with the medium harvested from control or FTO-depleted HepG2 cells and immunodepleted with either IgG or anti-DKK1 antibody. Relative luciferase activities were measured after 16 hours of WNT stimulation (WR, WNT3a and R-spondin1). n = 3. (F) Antibody neutralization assay. Control or FTO-depleted HeLa cells were transiently transfected with WNT reporter (TOP-flash). Cells were treated with either mock or WNT3A and R-spondin1 (WR), together with IgG or anti-DKK1 antibody in the culture medium (see methods for details). Relative luciferase activities were measured. n = 3. (G) Fragmented poly(A)+ RNAs from control or FTO-depleted HeLa cells were immunoprecipitated with anti-m6A antibody. Primer sets targeting different regions in the DKK1 gene (A: 5’UTR, B: CDS, C: 3’UTR targeting) were used for RT-qPCR. m6A enrichment were calculated as relative amounts of m6A immunoprecipitated fraction compared with input. Fold changes were compared after GAPDH normalization between input samples of control and FTO depletion. n = 3. (H) DKK1 RNA levels were measured in control or FTO-depleted SW480 cells treated with actinomycin D for the indicated time. GAPDH was used as a normalization control. n = 2. (I) RNA levels of mature and pre-spliced form of DKK1 (pre-DKK1) were measured by RT-qPCR from shCon or two different shFTO (#2 and #3) expressing HCT-8 cells. GAPDH was used as a normalization control. n = 3. (J) RNA levels of mature and pre-spliced form of DKK1 were measured by RT-qPCR from control or FTO-depleted HeLa cells (shFTO#3). GAPDH was used as a normalization control. n = 3. (K) RNA levels of mature DKK1 or pre-DKK1 were measured in control or FTO-depleted HepG2 cells (shFTO#2) by RT-qPCR. GAPDH was used as a normalization control. n = 2. Error bars are ± s.e.m. One-sided t-test. **: p<0.01, *: p<0.05.

**FIGURE 4-figure supplement 1.**
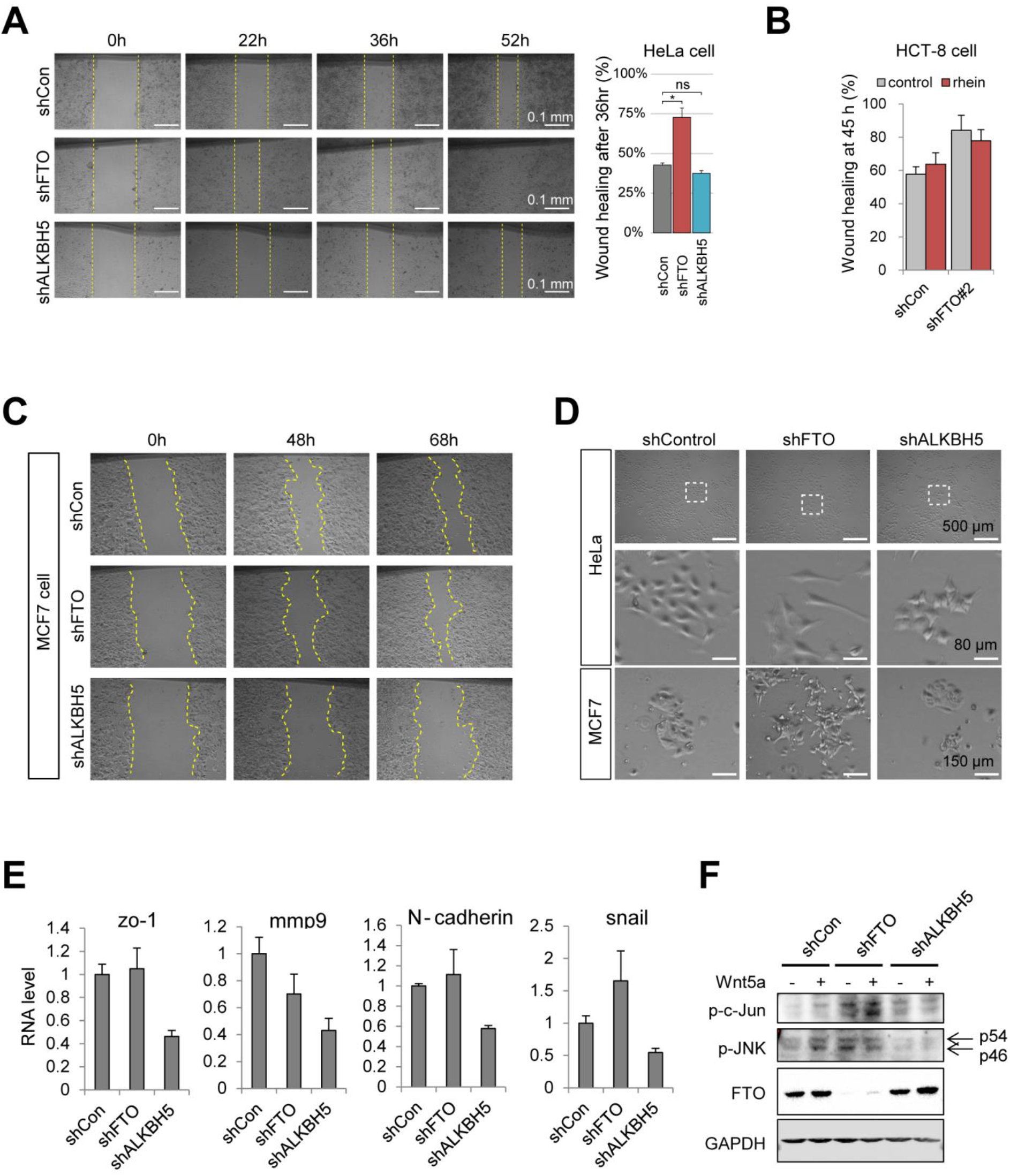
Loss of FTO promotes cell migration and polarity via WNT/PCP activation. (A) Wound healing assays in control, FTO- or ALKBH5-depleted HeLa cells. Wound widths at 0, 22, 36 and 52 hours after scratch were measured using imageJ software. Representative results are shown in left. Yellow dashed lines mark wound edges. Wound healing % at 36 hours after scratch were graphed (right). n = 3. (B) Representative wound healing activity at 45 hours after scratch in control or FTO-depleted HCT-8 cells (shFTO#2) treated either with DMSO control or rhein. n = 4. (C) Representative wound healing assay in control, FTO- or ALKBH5-depleted MCF7 cells. Yellow dashed lines indicate wound edges. (D) Bright field cell images of control, FTO- or ALKBH5-depleted HeLa and MCF7 cells with different magnification (see scale bars). (E) RNA levels of various EMT (epithelial mesenchymal transition) markers were measured by RT-qPCR from control, FTO- or ALKBH5-depleted HeLa cells. n = 3. (F) Control, FTO- or ALKBH5-depleted HeLa cells were stimulated with either mock (-) or with recombinant WNT5a (+). Protein levels of phosphorylated c-JUN (p-c-Jun), phosphorylated JNK (p-JNK) and FTO were measured by western blot. GAPDH was used as a loading control. Error bars are ± s.e.m. One-sided t-test. **: p<0.01, *: p<0.05, n.s.: p>0.05. shFTO#3 was used for FTO depletion.

**FIGURE 5-figure supplement 1.**
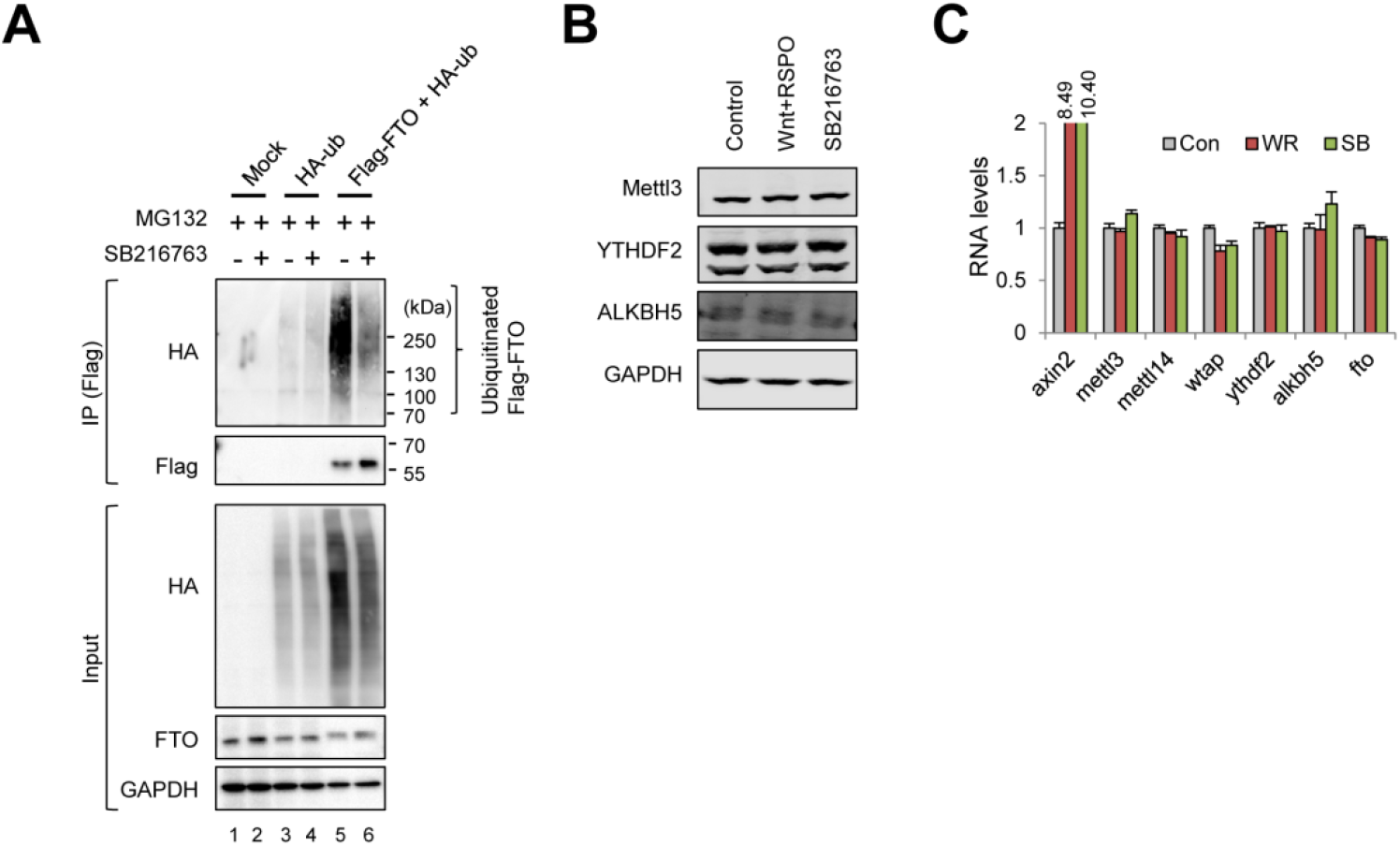
WNT/STOP signaling attenuates FTO protein polyuniquitination and degradation. (A) Polyubiquitination of Flag-FTO in the presence of proteasomal inhibitor (MG132) and GSK3 inhibitor (SB216763). HeLa cells were transiently transfected with either mock, HA-ub alone or Flag-FTO together with HA-ub. After Flag immunoprecipitation, ubiquitinated FTO were assessed by western blot using HA antibody. (B) Protein levels of m6A pathway writer (METTL3), reader (YTHDF2) or eraser (ALKBH5) were measured by western blot from HeLa cells stimulated with WNT (WNT+RSPO, WNT3a and R-spondin1) or GSK3 inhibitor SB216763. GAPDH was used as a loading control. (C) RNA levels of various m6A pathway components were assessed by RT-qPCR from HeLa cells after WNT stimulation (WR, WNT3a and R-spondin1) or after GSK3 inhibition (SB) (n=3 each). Axin2 was used as a positive control for WNT activation. GAPDH was used as a normalization control. n = 3.

**FIGURE 6-figure supplement 1.**
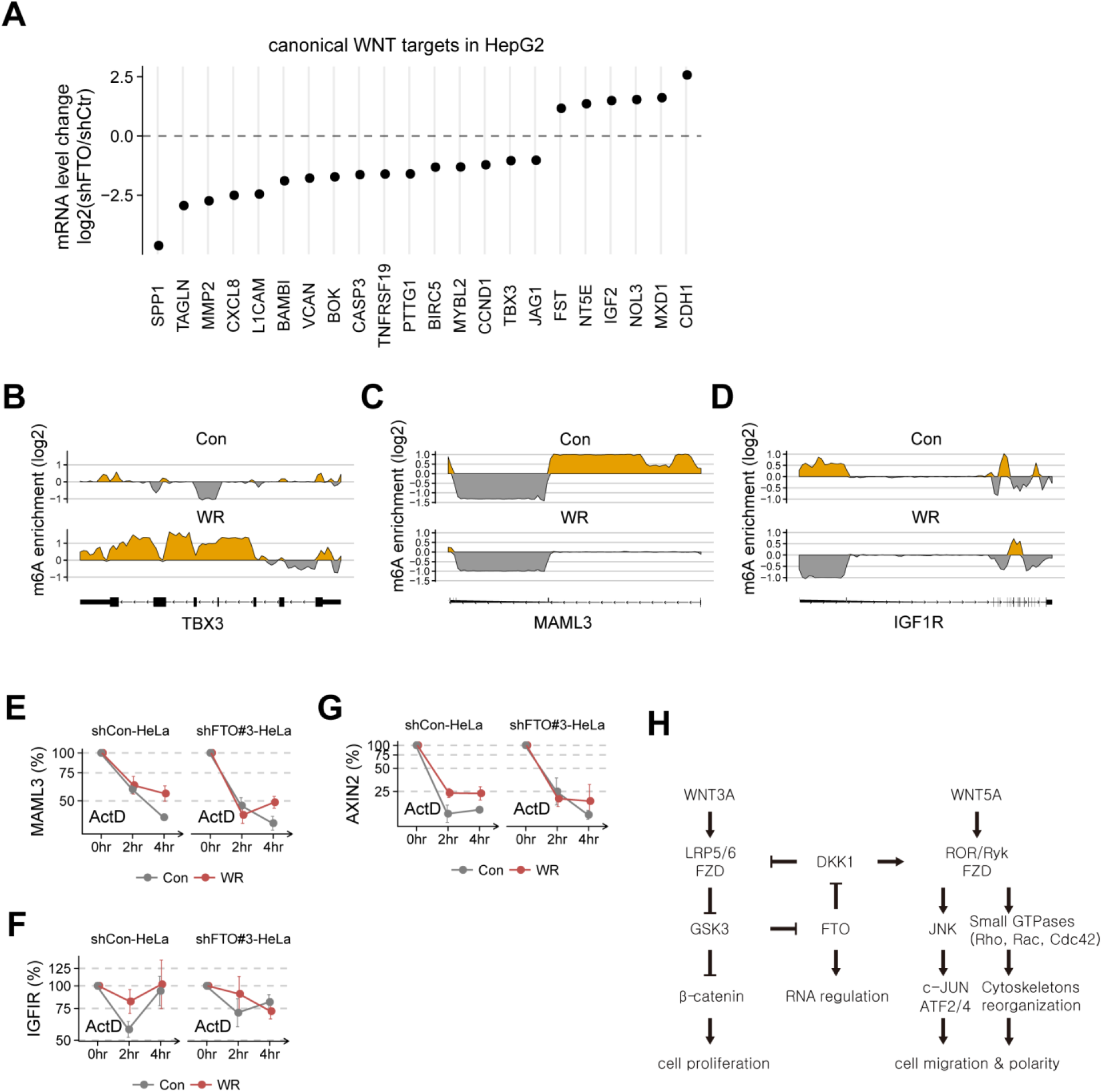
WNT signaling stabilizes FTO target mRNAs. (A) Regularized fold change of canonical WNT targets in FTO-depleted HepG2 cells. Only differentially expressed targets (adjusted p-value < 0.01) are shown. (B-D) Change in m6A enrichment (IP/Input) across the TBX3 (B), MAML3 (C) and IGF1R (D) genes after WNT3A treatment (WR). Read coverages of both IP and input samples were standardized as described in Methods. The yellow areas indicate putative m6A peaks. The gene models are shown below. (E-G) RNA levels of MAML3 (E), IGF1R (F) and AXIN2 (G) were measured by RT-qPCR in control or FTO-depleted HeLa cells treated with WNT (WR, WNT3A and R-spondin1) and actinomycin D (act D) for the indicated time (n = 3). GAPDH was used as a normalization control. (H) Our proposed model for the intricate interaction between FTO and multiple WNT pathways. WNT3A-induced canonical signaling pathway leads to the β-Catenin stabilization and cell proliferation while WNT5A-induced WNT/PCP pathway activates small GTPases such as Rho and Rac (not shown) and JNK to regulate cell migration and polarity. Loss of FTO upregulates DKK1 and inhibits canonical WNT/β-Catenin while activates non-canonical WNT/PCP pathways. Canonical WNT/STOP signaling stabilizes FTO protein and regulates mRNA stability in a FTO-dependent manner. Error bars are ± s.e.m.

## SOURCE DATA LEGENDS

Figure 2-source data 1. Uncropped gel images for Figure 2.

Figure 3-source data 1. Uncropped gel images for Figure 3B.

Figure 4-source data 1. Uncropped gel images for Figure 4E.

Figure 5-source data 1. Uncropped gel images for Figure 5.

Figure 2-figure supplement 1-source data 1. Uncropped gel images for Figure 2-figure supplement 1.

Figure 3-figure supplement 1-source data 1. Uncropped gel images for Figure 3-figure supplement 1C-D.

Figure 4-figure supplement 1-source data 1. Uncropped gel images for Figure 4-figure supplement 1F.

Figure 5-figure supplement 1-source data 1. Uncropped gel images for Figure 5-figure supplement 1A.

Figure 5-figure supplement 1-source data 2. Uncropped gel images for Figure 5-figure supplement 1B.

